# Effective Ultrasonic Stimulation in Human Peripheral Nervous System

**DOI:** 10.1101/2021.04.22.440931

**Authors:** Thomas Riis, Jan Kubanek

## Abstract

**Objective:** Low-intensity ultrasound can stimulate excitable cells in a noninvasive and targeted manner, but which parameters are effective has remained elusive. This question has been difficult to answer because differences in transducers and parameters—frequency in particular—lead to profound differences in the stimulated tissue volumes. The objective of this study is to control for these differences and evaluate which ultrasound parameters are effective in stimulating excitable cells.

**Methods:** Here, we stimulated the human peripheral nervous system using a single transducer operating in a range of frequencies, and matched the stimulated volumes with an acoustic aperture.

**Results:** We found that low frequencies (300 kHz) are substantially more effective in generating tactile and nociceptive responses in humans compared to high frequencies (900 kHz). The strong effect of ultrasound frequency was observed for all pressures tested, for continuous and pulsed stimuli, and for tactile and nociceptive responses.

**Conclusion:** This prominent effect may be explained by a mechanical force associated with ultrasound. The effect is not due to heating, which would be weaker at the low frequency.

**Significance:** This controlled study reveals that ultrasonic stimulation of excitable cells is stronger at lower frequencies, which guides the choice of transducer hardware for effective ultrasonic stimulation of the peripheral nervous system in humans.

## I. Introduction

Low-intensity focused ultrasound has the potential to transform diagnoses and treatments of nervous system disorders. Ultrasound can be remotely and flexibly focused into a specific target, where it modulates the activity of excitable cells and nerve fibers transiently for brief stimuli [1], [2], and induces plastic reorganization for longer stimuli [3]–[9]. In the common range of neuromodulatory frequencies, 0.25–1.0 MHz, ultrasound combines sharp focus with exquisite depth penetration, including through the intact human skull [10]–[14]. Ultrasound has thus become the only neuromodulation modality that offers the full triad of desirable properties—noninvasiveness, focus, and depth penetration.

Despite several decades of research efforts [3], [15]–[22], it is not known which ultrasound parameters should researchers and clinicians use to stimulate excitable cells and nerves effectively. The ultrasound frequency is a key parameter: it governs the choice of the transducer to be used, the size of the focal region, and depth penetration. Yet, whether lower [20]–[23] or higher [24], [25] frequencies stimulate the nervous system more effectively is unknown. This question could not be addressed in previous studies because changes in frequency lead to large changes in the stimulated volumes, and therefore to large changes in the probability of registering a response [24]. For example, the focal volume of a 300 kHz focused ultrasonic wave, compared to 900 kHz of otherwise equal parameters, is 27 times larger. The stimulation volume has therefore presented a major confound in previous studies and, consequently, the effect of stimulation frequency remains unknown.

Here, we used a single ultrasonic transducer and matched stimulation volumes for different frequencies using a metallic aperture. We applied the stimuli to the peripheral nervous system in humans, receptors and nerve endings of the index finger in particular. The stimulation of the peripheral nervous system has two main advantages for the investigation of the effect of frequency. First, the approach provides an acoustically transparent access to excitable cells and nerve fibers [17], [23], [26]. This is in contrast to a transcranial application, which would lead to substantial relative attenuation of the high frequency. Second, the approach does not suffer from auditory and vestibular artifacts that confound applications of ultrasound through the skull [27]–[29].

In peripheral nerves, ultrasound of frequencies between 0.25 MHz and 7 MHz can decrease [30]–[36] or increase [37]–[40] specific features of action potential conduction. When of sufficient pressure, ultrasound can also directly trigger action potentials [41]–[46]. How the outcome depends on frequency has been difficult to determine because effects of frequency have not been compared within each study and within particular experimental condition. In peripheral nerve structures and endings, relatively low ultrasound pressures in a frequency range between 0.48 MHz and 2.67 MHz can elicit tactile [26], [47]–[50], nociceptive [26], [48], [51]–[53], and auditory [49], [54], [55] responses, depending on the stimulated structure. Tactile and nociceptive responses appear to be more effectively triggered at lower frequencies [23], but the effects have been confounded by the unmatched stimulated volumes.

We tested the effects of ultrasound at 300 kHz and 900 kHz using the same transducer and matched the stimulated volumes. We varied frequency along with 6 other parameters. Across ultrasound parameters and subjects’ responses, we arrived at a common and salient finding: lower frequencies are more effective in stimulating mechanoreceptors and nerve endings. This finding guides the choice of ultrasound frequency for modulations of the peripheral nervous system, and provides new information for theories of ultrasound interactions with biological tissues.

## II. Methods

### A. Subjects and apparatus

Twenty-six subjects (19 males, 7 females, aged between 20-37 years) participated in this study. Data of all subjects were included in the analyses; no subject was excluded. The study was approved by the Institutional Review Board of the University of Utah. Subjects were asked to gently rest the finger on a metallic aperture under a 45-degree angle. The aperture and the participants’ finger were immersed in water (Fig. 1a). The aperture (2 mm-thick aluminum) had a 4 mm in diameter opening for the ultrasound beam. The height of the case that held the aperture was 52 mm and its total diameter 70 mm.

**Fig. 1.**
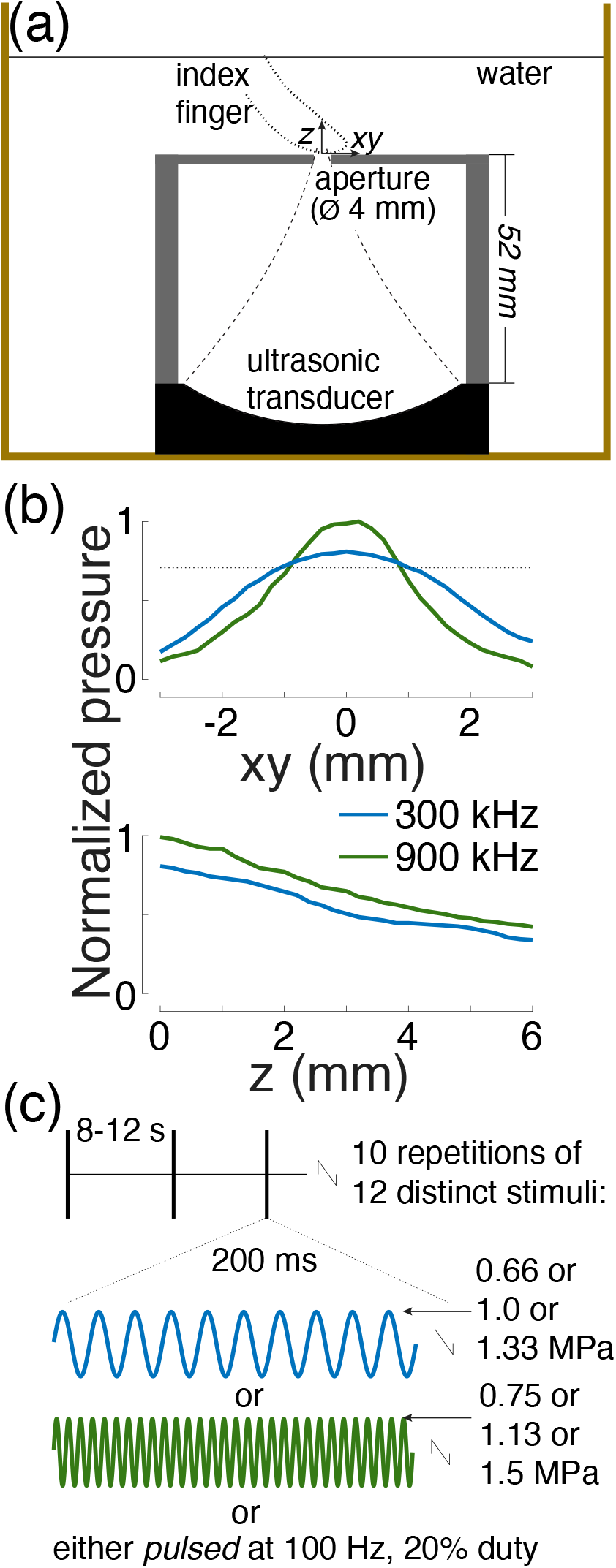
Ultrasound stimulation apparatus and stimuli. (a) Setup. A focused ultrasound transducer delivered a 300 kHz or 900 kHz stimulus into a subject’s index finger through water. The ultrasound beam profile (dashed lines) was controlled through an aperture 4 mm in diameter. The water was either one-time degassed (18 subjects) or continuously degassed (8 subjects), and was kept at about 30°C. (b) Peak-normalized ultrasound pressure field for the two frequencies. The aperture size and pressure levels were set such that both frequencies produced a comparable focal volume (see Methods). The pressure profile was averaged over the x and y dimensions. The dotted lines show the 0.707 (0.5) pressure (intensity) levels to characterize the fields using full-width-at-half-maximum values. (c) Stimuli. Each subject experienced 10 repetitions of 12 distinct stimuli. The stimuli, 200 ms in duration, were selected randomly and delivered each 8-12 seconds. We tested 2 frequencies, 3 pressure levels, and continuous and pulsed (100 Hz, 20% duty) stimuli.

The temperature of the water was maintained at around 30°C using a heater with temperature controller (HG-802, Hygger, Shenzhen, China). The controller continually monitored the bath temperature and turned on when temperature dropped below the target value. Eighteen subjects performed the experiment in one-time degassed water (“still water”). Eight subjects performed the experiment within a continuously degassed water tank system (AIMS III system with AQUAS-10 Water Conditioner, Onda, Sunnyvale, USA). The water conditioner (AQUAS-10) treats water for ultrasound measurements in compliance with IEC 62781. The conditioner degasses water to remove undesired bubbles, removes suspended particles and biological contaminants, and deionizes water. The dissolved oxygen is between 2.0-2.5 PPM during continuous operation, according to measurements provided by the manufacturer (Onda). In comparison, tap water contains about 10.5 PPM of dissolved oxygen.

A focused ultrasonic transducer (H-115, Sonic Concepts, 64 mm diameter, 52 mm focal depth) was positioned 52 mm below the aperture (Fig. 1a). The transducer was operated at 300 kHz and the third harmonic, 900 kHz. Stimuli were generated by a custom Matlab program that produced the stimulation waveforms in a programmable function generator (33520b, Keysight, Santa Rosa, USA). The signals were amplified using a 55-dB, 300 kHz–30 MHz power amplifier (A150, Electronics & Innovation, Rochester, USA).

Subjects had their eyes closed and wore noise-cancelling earmuffs (X4A, 3M, Saint Paul, USA; noise reduction rating of 27 dB) so that they could fully focus on the ultrasonic stimuli. Subjects could not hear or see the stimuli or their generation.

### B. Stimuli

We used 12 distinct stimuli, of all combinations of 2 frequencies, 3 pressure levels, and 2 stimulus kinds. The parameters were chosen to provide safe and effective stimulation. The two carrier frequencies (300 kHz and 900 kHz) were chosen such that the ultrasound could penetrate into depth with minimal attenuation, yet could be focused into focal regions of dimensions on the order of millimeters. The duration of each stimulus (200 ms) was chosen to be long enough to provide effective stimulation yet short enough to enable repeated stimulation in compliance with the FDA guidelines (see Stimulus safety). In particular, previous studies in the peripheral and central nervous system demonstrated that the stimulatory effects of ultrasound increase with stimulus duration, and the effects saturate at around 100-200 ms [25], [47], [51], [56]. The majority of previous studies used pulses that were shorter than 100 ms [17], [26], [34], [36], [41], [43], [45], [46], [52], [53], and so effective stimulation could likely also be achieved with stimuli on the order of milliseconds. The peak pressure amplitudes measured at the center of the aperture were 0.66 MPa, 1.0 MPa, and 1.33 MPa for 300 kHz, and 0.75 MPa, 1.13 MPa, and 1.5 MPa for 900 kHz. The peak pressures were chosen such as to comply with the *I*_SPPA_ Track 3 510(k) [57] recommendation for each pulse and within the *I*_SPPA_ recommendation over the course of the experiment (see Stimulus safety). These pressure levels provided robust response rates. The stimuli were either continuous (200 ms of tone burst) or pulsed at 100 Hz at 20% duty (i.e., 2 ms on, 8 ms off, 2 ms on, etc., for 200 ms). The continuous and pulsed stimuli had the same pressure amplitude. The pulse repetition frequency was chosen to match the frequency responsiveness of mechanoreceptors [25], [58]. The duty cycle was set low enough to provide sufficient contrast for distinguishing mechanical from thermal effects.

The pressure fields were measured using a capsule hydrophone (HGL-0200, Onda) calibrated between 250 kHz and 40 MHz and secured to 3-degree-of-freedom programmable translation system (Aims III, Onda). There were 10 repetitions of the 12 stimuli, producing a total of 120 stimulation trials per subject. The stimuli were delivered every 8-12 s. The stimuli were drawn from the 120-stimulus set randomly without replacement. This way, stimulus order could not affect the results.

The stimuli used in the study had most of their energy concentrated at the carrier frequency or in its near proximity. The 300 and 900 kHz continuous stimuli had a −20-dB pressure amplitude bandwidth of 31Hz and 27 Hz, respectively. The 300 and 900 kHz pulsed stimuli both had a −20-dB pressure amplitude bandwidth of 2.6 kHz

### C. Responses and their assessment

Subjects were instructed to report a percept with a verbal command of either {Pain, Warm, Cold, Vibration, Tap}. The respective capital letters were noted by the experimenter onto a sheet, into the appropriate stimulus bracket made explicit by the running Matlab program. Following the experiment, for each stimulus type, response frequency was computed as the proportion of trials in which a response was registered, relative to the 10 repetitions. In a small proportion (<1%) of cases, subjects reported two percepts; the first one was registered.

### D. Aperture choice

The aperture was made inside a 2-mm layer of aluminum. The acoustic impedance of this metal is 17.1 MRayl [59]. Given the double reflection from the two interfaces (water-metal, metal-water), only 8.6% of the energy could reach the opposite side of the metallic layer. The aperture therefore served as an effective spatial filter for the ultrasound beam. The aperture diameter was chosen to be large enough so that a 300 kHz beam could effectively penetrate the soft tissues, but small enough so that the 300 kHz beam was comparable to the 900 kHz beam. Due to diffraction effects at the edges of the metal, the 300 kHz and 900 kHz beams could be matched satisfactorily but not exactly (Fig. 1). To compensate for the differences, the pressure amplitudes of the 900 kHz beam were scaled up by 1 dB compared to 300 kHz. This adjustment brought the focal volumes of the two beams close to each other (within 10%; see Results). The gradients of the 300 kHz and 900 kHz fields were comparable and relatively omni-directional.

### E. Acoustic continuum

The acoustic impedance of water and skin, including soft tissues, are closely matched (1.48 MRayl compared to 1.68 MRayl [60]) This way, about 99.6% of the energy, 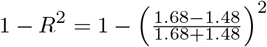, was delivered into the finger. The water-finger interface is therefore essentially acoustically transparent and can be considered as a continuum from the perspective of neuromodulatory ultrasound.

### F. Stimulus safety

The ultrasonic stimuli used in this study were below the FDA 510(k) Track 3 recommendations [57]. In particular, the highest peak pressure used in the study, 1.5 MPa, corresponds to peak intensity of 67.0 W/cm^2^, which is well below the FDA recommendation of *I*_SPPA_ = 190 W/cm^2^. In addition, the time-average spatial peak intensity was *I*_SPTA_ = 430 mW/cm^2^, also below the FDA recommendation of *I*_SPTA_ = 720 mW/cm^2^. The *I*_SPPA_ and *I*_SPTA_ values for the individual stimuli are provided in Table II.

## III. Results

We applied 12 distinct ultrasound stimuli, 10 repetitions each, to the index finger of 26 humans (Fig. 1). The ultrasound was delivered into the tissues in water (Fig. 1a).

We used a focused ultrasonic transducer (Methods) that operated at 300 kHz or 900 kHz. Without any control, the focal volume of the 300 kHz beam would be 27 times larger than that of 900 kHz, since both the width and the focal length of the beam are proportional to wavelength [61]. To match the stimulated volumes, we constrained the beam using a metallic aperture with an opening set such that both frequencies produced comparable stimulation volumes (Fig. 1b; see Methods). Indeed, the full-width-at-half-maximum (FWHM) volumes were 4.4 and 4.8 mm^3^ for the 300 kHz and 900 kHz beams, respectively. The FWHM diameter was 2.0 mm and 1.6 mm, and FWHM focal length 1.4 mm and 2.4 mm for the 300 kHz and 900 kHz beams, respectively. The individual stimuli (Fig. 1c) were delivered randomly every 8-12 seconds.

We found that the low frequency was more than twice as effective in eliciting subjects’ responses compared to the high frequency (Fig. 2). Specifically, subjects responded to the 300 kHz stimuli on average in 41% of cases compared to just 20% of cases for the 900 kHz stimuli. This difference was highly significant (*p* = 6.5 × 10^−8^, paired t-test, *t*_25_ = 7.6). This finding may appear surprising given that the 900 kHz stimulus, if anything, impacted a slightly larger focal volume and produced slightly higher peak pressures (Fig. 1b).

**Fig. 2.**
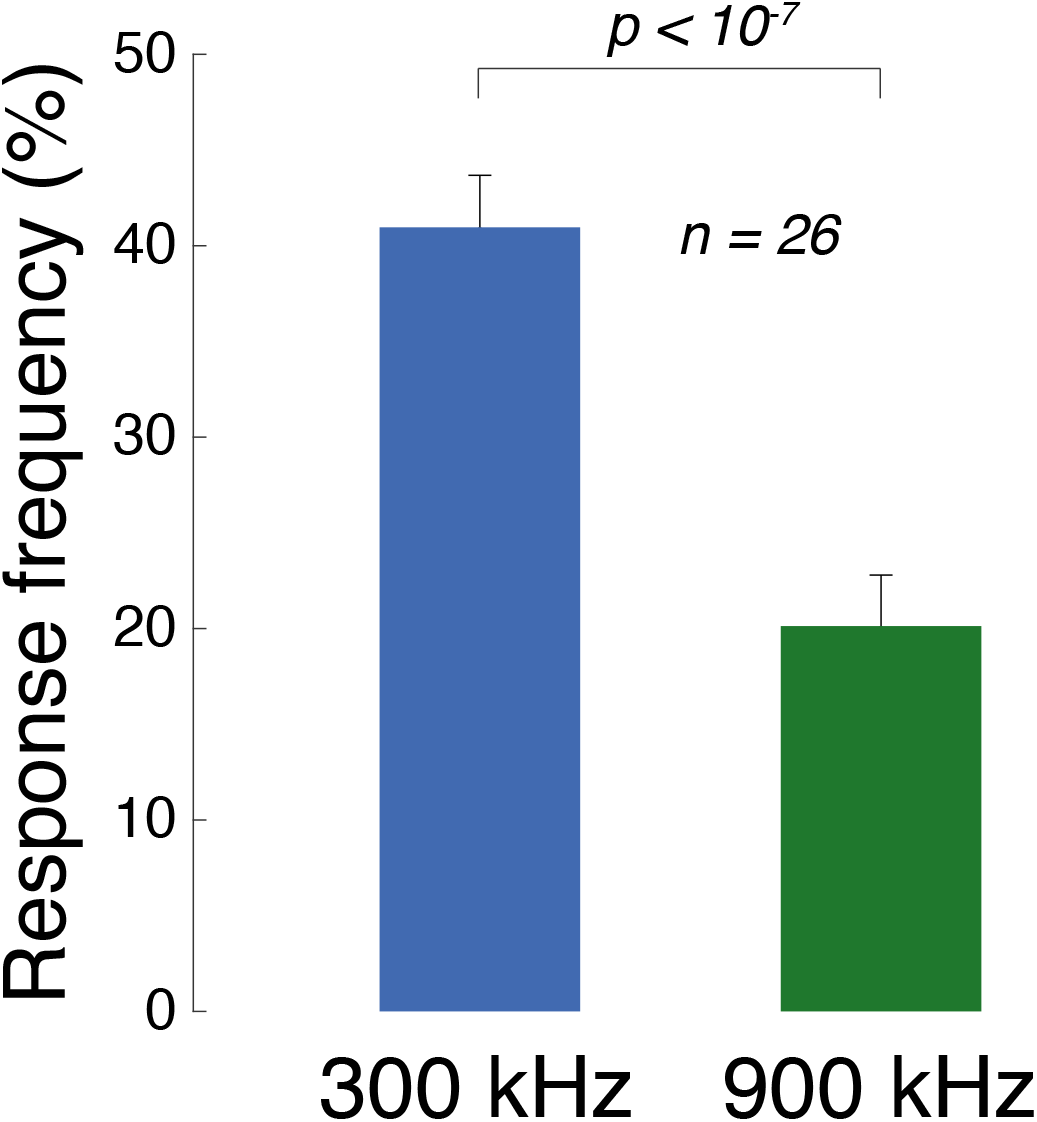
Low frequencies stimulate excitable cells more effectively than high frequencies. Mean±s.e.m. response frequency for all 300 kHz stimuli (blue) and all 900 kHz stimuli (green). The data contain responses from all 26 subjects. Significance was assessed using paired t-test.

We next asked whether this prominent effect of ultrasound frequency depends on the ultrasound pressure magnitude, type of stimulus, or the subjects’ responses. Across these conditions, the lower frequency was consistently found to be more effective (Fig. 3). We assessed the significance of these effects using a 3-way ANOVA, with factors frequency (*F*), pressure (*P*), stimulus kind (*S*; continuous or pulsed), and all possible interactions between these factors (Table I). This linear model confirms that frequency *F* is a highly significant main factor, even when including all possible interactions. Notably, Fig. 3-top reveals that the modulation of the response frequency by pressure is stronger for the low frequency. This is statistically confirmed in Table I, which detected a highly significant *F* × *P* interaction. This is an important finding because within a given frequency, the ultrasound beam geometry is fixed. Therefore, the lower frequency produces stronger effects even when the beam geometry is fixed. In addition, the contrast between the pulsed and the continuous stimuli is more pronounced at the lower frequency (Fig. 3-middle). This is confirmed by a significant *F* × *S* interaction. The higher effectiveness of the lower frequency is observed also when we separately focus on tactile and nociceptive responses (Fig. 3-bottom), and when we build separate ANOVAs for these two response kinds (Table I, middle and right columns).

**Figure 3.**
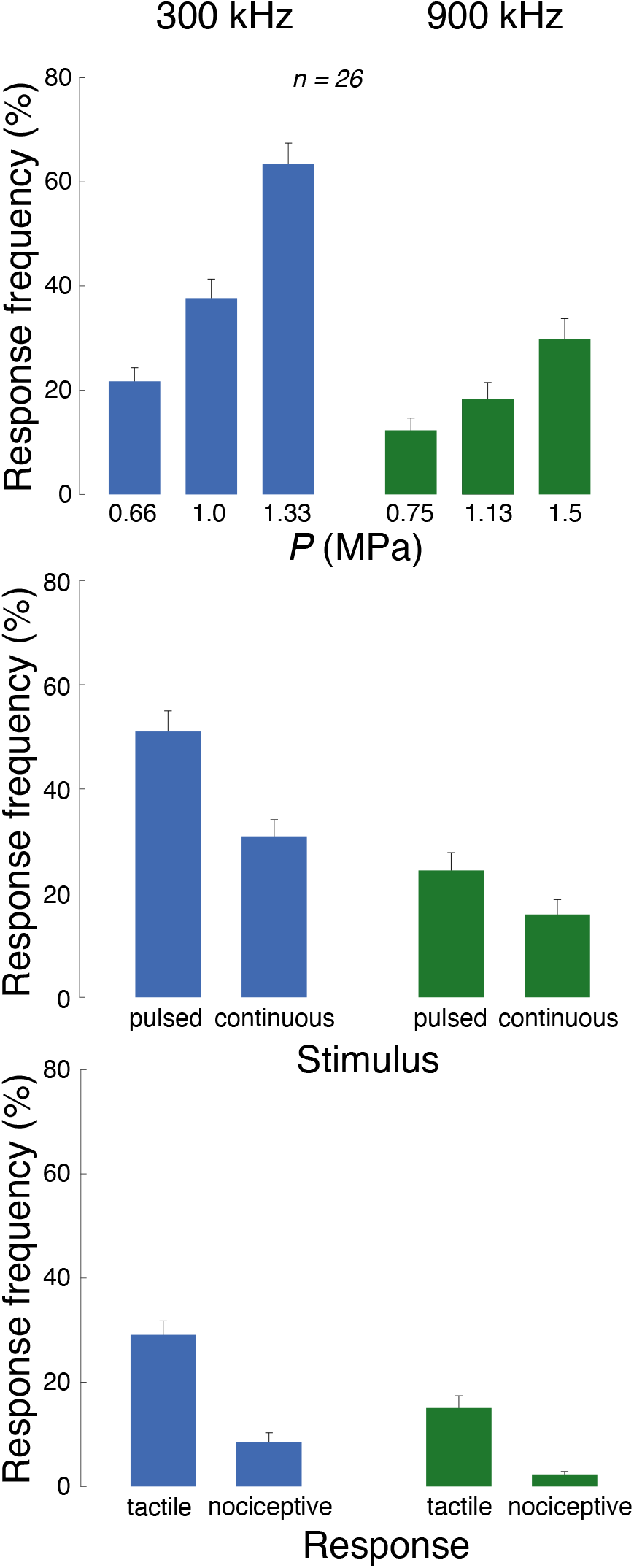
Low frequencies stimulate excitable cells more effectively than high frequencies, in all tested conditions. Mean±s.e.m. response frequency for the specific stimuli at the 300 kHz carrier frequency (blue) and 900 kHz (green). Response frequency is presented as a function of the ultrasound pressure amplitude (top row), stimulus kind (continuous or pulsed; middle row), and the type of subjects’ response (tactile (tap or vibration) or nociceptive; bottom row). The significance of these effects is assessed using an omnibus linear model (Table I).

**TABLE I.** Effects of stimulus parameters. The effects of ultrasound frequency (***F***), pressure (***P***), and stimulus kind (***S***; continuous or pulsed) on the response frequency (any response; left column) and specifically on tactile (middle column) and nociceptive responses (right column). These effects were assessed using a three-way ANOVA model that features these three main effects and all possible interactions. Bold entries are significant (***p*** < **0.05**).

|  | Any response | Tactile | Nociceptive |
| --- | --- | --- | --- |
| $F$ | $7.4 \times 10^{-15}$ | $1.2 \times 10^{-8}$ | $4.0 \times 10^{-6}$ |
| $P$ | $3.2 \times 10^{-18}$ | $2.6 \times 10^{-11}$ | $1.5 \times 10^{-8}$ |
| $S$ | $4.3 \times 10^{-8}$ | $4.3 \times 10^{-15}$ | **0.0019** |
| $F \times P$ | **0.00058** | 0.097 | **0.00048** |
| $F \times S$ | **0.022** | **0.0001** | 0.079 |
| $P \times S$ | **0.0078** | $7.2 \times 10^{-5}$ | **0.046** |
| $F \times P \times S$ | 0.71 | 0.27 | 0.21 |

The data for all stimulus conditions are presented separately for the individual responses in Fig. 4. The figure supports the findings of Fig. 3 and Table I that the lower frequency is more effective, regardless of response kind. With respect to nociceptive responses, these data replicate previously reported response rates [26], [51] and show that lower frequencies increase the likelihood of registering a nociceptive response.

**Fig. 4.**
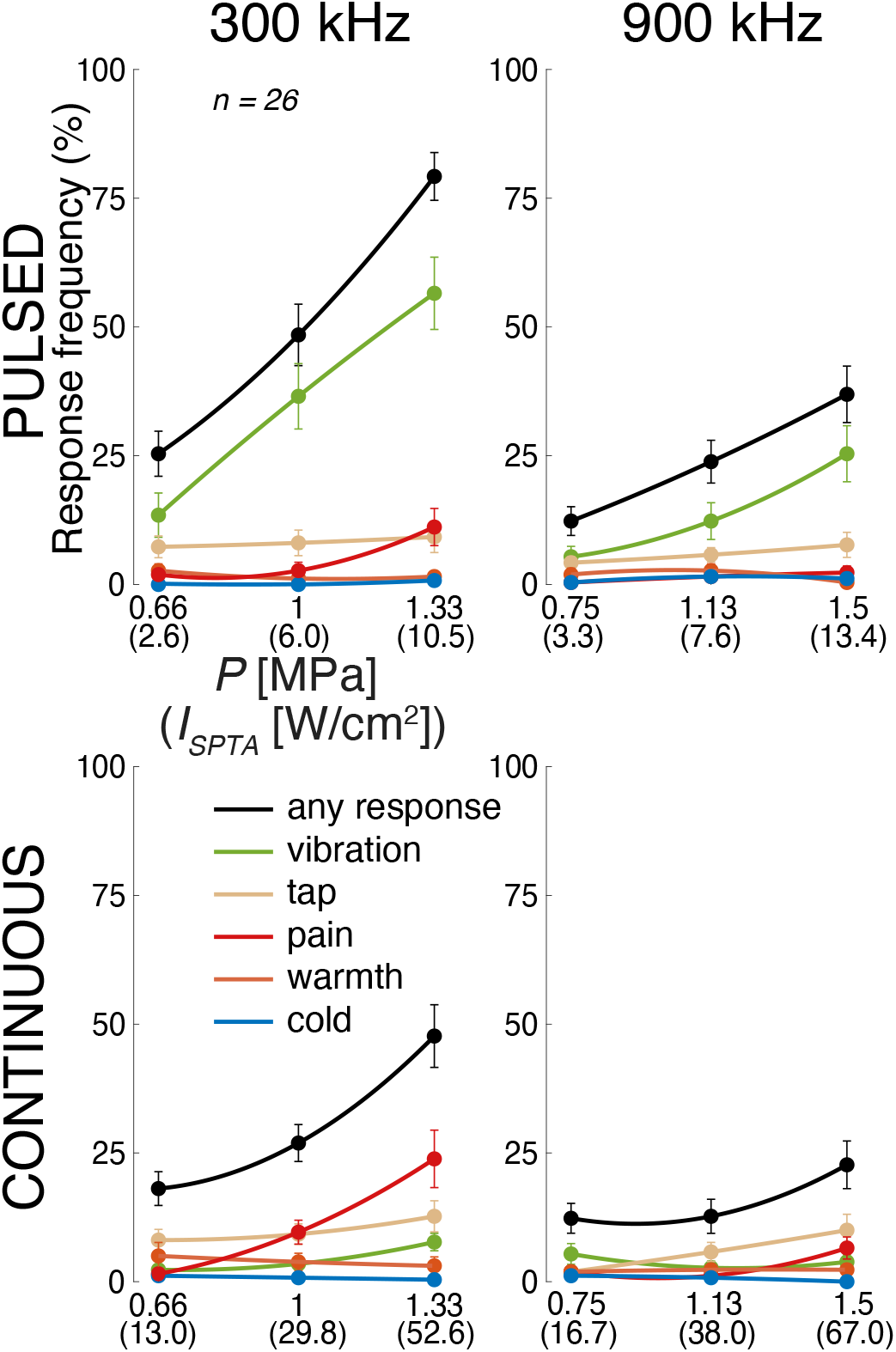
Individual responses to the individual stimuli. Mean±s.e.m. response frequency separately for each of the 12 stimuli, and separately for the individual response kinds (see legend). The abscissa provides peak pressure (***I***_SPTA_) values. The lines represent second-order polynomial fits.

These data reveal an additional salient finding. Tactile (vibration in particular; green) responses are more frequently observed for the pulsed stimuli (top row), whereas nociceptive stimuli (red) are more frequently observed for the continuous stimuli (bottom row). This phenomenon is analyzed in a dedicated Fig. 5. Pulsed (continuous) stimuli elicited a mean tactile response frequency of 44% (14%), and this difference is significant (*p* =1.8 × 10^−6^, paired t-test, *t*_25_ = −6.2). Reversely, pulsed (continuous) stimuli elicit a mean nociceptive response frequency of 5% (12%), and this difference is significant (*p* = 0.0025, paired t-test, *t*_25_ = 3.4). Thus, there is a double dissociation of the response kind through the stimulus kind. This finding, combined with the qualitatively distinct natures of the tactile and painful responses reported by the subjects, suggests that the distinct kinds of stimuli activated distinct receptors in the finger. This notion is elaborated on in the Discussion.

**Fig. 5.**
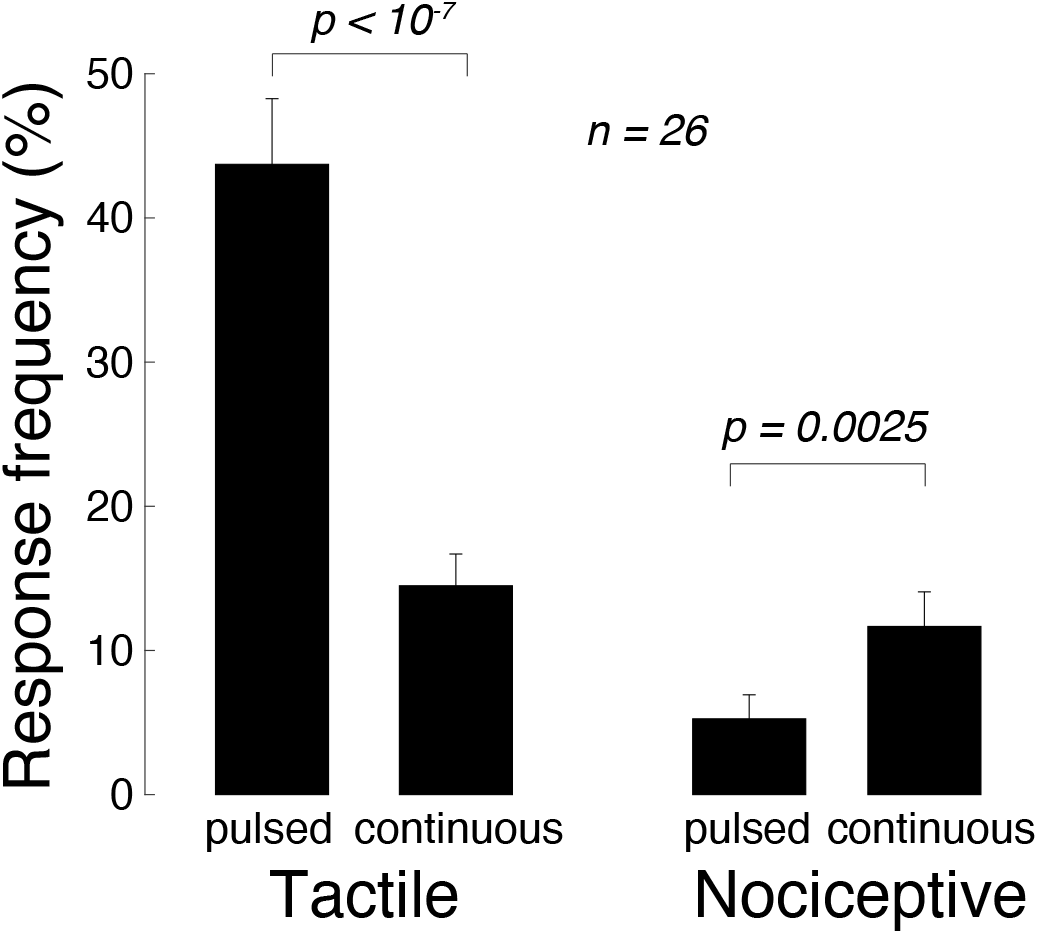
Pulsed (continuous) stimuli more effectively elicit tactile (nociceptive) responses. Mean±s.e.m. response frequency for vibrotactile (left) and nociceptive (right) responses, separately for pulsed and continuous stimuli. Significance is assessed using paired t-tests.

The finding that nociceptive responses are more likely to be elicited by continuous than pulsed stimuli suggests an involvement of a mechanism that delivers a greater amount of total energy into the target, such as heating. However, if the effect was due to heating, the effect should increase with frequency (see Discussion). Yet, effects were stronger for the lower frequency compared to the higher frequency also for these continuous stimuli (Fig. 4; 12% vs 3%; *p* = 0.0027, paired t-test, *t*_25_ = 3.3).

We tested whether lower frequencies are more effective also when there is no direct contact of the finger with the volume-controlling aperture. In this regard, we recorded additional responses in one subject who used the thumb and placed this finger approximately 1 mm above the aperture, in degassed water. The response frequencies in this subject were 65% and 33% for the 300 kHz and 900 kHz stimuli, respectively. This replicated the subject’s index finger effects of 55% and 13%, respectively. Thus, no contact with the aperture was necessary to elicit these effects [23].

The stimuli used in this study complied with the FDA 510(k) Track 3 recommendations [57] (see Methods and Table II). The stimulation effects were transient. No signs of harm were detected by the experimenter or reported by the subjects. Finger sensation was normal following the data collection. In one subject, the skin at the stimulated zone appeared redder. The color was back to normal within several hours. The subject noted to have irritable skin and so the effect could have been due to the prolonged contact of the finger with the metallic aperture.

**TABLE II.**
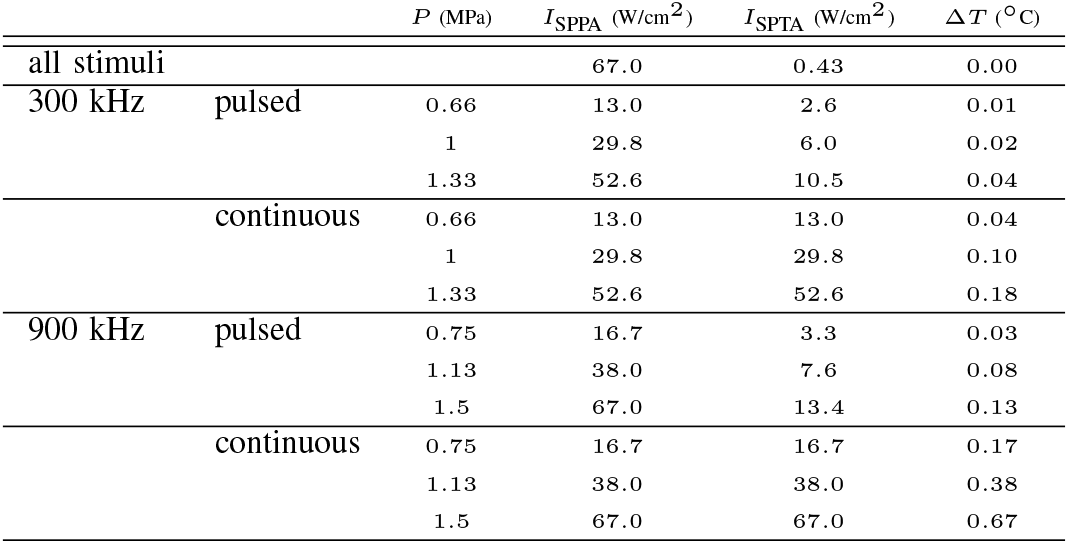
Stimulus levels. The peak intensity of each stimulus ***I***_SPPA_ was computed as 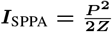, where the acoustic impedance of the skin ***Z*** = **1.68** MRayl [60]. The time average intensities ***I***_SPTA_ were ***I***_SPTA_ = ***I***_SPPA_ for continuous stimuli and ***I***_SPTA_ = **0.2*****I***_SPPA_ for the pulsed stimuli (20% duty cycle). The first row provides ***I***_SPTA_ over the course of the experiment. This value takes into account the 10 s inter-stimulus period, i.e., mean(***I***_SPTA_)×**0.2/10**. The maximal temperature rise values for the individual 0.2 s stimuli were computed as in [51] using the same attenuation coefficient (0.1 Np/cm/MHz).

| | | $P$ (MPa) | $I_{SPPA}$ (W/cm<sup>2</sup>) | $I_{SPTA}$ (W/cm<sup>2</sup>) | $\Delta T$ (°C) |
| --- | --- | --- | --- | --- | --- |
| all stimuli |  |  | 67.0 | 0.43 | 0.00 |
| 300 kHz | pulsed | 0.66 | 13.0 | 2.6 | 0.01 |
|  |  | 1 | 29.8 | 6.0 | 0.02 |
|  |  | 1.33 | 52.6 | 10.5 | 0.04 |
|  | continuous | 0.66 | 13.0 | 13.0 | 0.04 |
|  |  | 1 | 29.8 | 29.8 | 0.10 |
|  |  | 1.33 | 52.6 | 52.6 | 0.18 |
| 900 kHz | pulsed | 0.75 | 16.7 | 3.3 | 0.03 |
|  |  | 1.13 | 38.0 | 7.6 | 0.08 |
|  |  | 1.5 | 67.0 | 13.4 | 0.13 |
|  | continuous | 0.75 | 16.7 | 16.7 | 0.17 |
|  |  | 1.13 | 38.0 | 38.0 | 0.38 |
|  |  | 1.5 | 67.0 | 67.0 | 0.67 |

## IV. DISCUSSION

In this study, we addressed the question whether low or high frequencies are more effective in stimulating excitable cells in the peripheral nervous system. We clamped the stimulated volumes and therefore prevented their large confounding effects. We harnessed the intact human peripheral nervous system that is acoustically transparent, is not influenced by auditory or vestibular artifacts, and preserves all mechanical ties between cells and tissues [62]. Operating in the common range of ultrasound stimulation frequencies, we found multiple lines of evidence supporting a conclusion that a low, 300 kHz frequency is more effective than a high, 900 kHz frequency (Fig. 2, Fig. 3, and Table I).

This result validates data of previous studies in the peripheral nervous system that suggested that lower frequencies may be more effective [23], [48], [49], [51]. Our volume-clamped experiment now draws a firm conclusion on this literature: lower frequencies are more effective.

From the translational standpoint, this result has one benefit and one drawback. Stimulation at lower frequencies minimizes ultrasound attenuation and thus facilitates the delivery of ultrasound into the desired excitable target. On the other hand, at 300 kHz the ultrasound wavelength is about 5 mm, which limits the spatial resolution of the approach to about that order.

Unexpectedly, we found a double dissociation of the response kind through the stimulus kind (Fig. 5). In particular, pulsed stimuli were more effective in eliciting vibrotactile responses than continuous stimuli. This effect was reversed for nociceptive responses—continuous stimuli were more effective than pulsed stimuli in eliciting nociceptive responses (Fig. 5). The qualitatively distinct kinds of subjects’ responses suggest that these distinct kinds of ultrasonic stimuli modulated distinct sets of excitable cells. The pulsed 100Hz stimuli may elicit forces that are modulated at 100 Hz, such as radiation force, which may activate Pacinian corpuscles and other mechanoreceptors with sensitivity in that range [47], [49]. The low-frequency- or steady-displacement-sensing Meissner corpuscles, Merkel cells, and Ruffini endings may also be partially implicated, especially in the less frequent tap sensations [58]. In comparison, nociception in the skin is mediated by free nerve endings. These neural pathways most commonly involve unmyelinated, small-diameter C-fiber axons, but can also constitute myelinated A-fiber axons [63].

The free nerve endings that mediate nociception in the skin express several classes of ion channels that have been shown to be activated by ultrasound. These include voltage-gated sodium channels [18], [64], K2P channels [64], TRPA1 channels [65], TRPC1 channels [66], ASIC channels [67], and Piezo channels [68]. Given our finding that low frequencies are more effective, even for the continuous stimuli, we can conclude that some of these channels respond to the mechanical, as opposed to thermal, aspects of the ultrasound. A mechanical activation of ion channels is becoming a predominant hypothesis of the biophysical action of ultrasound on excitable cells [1], [2], [10], [11], [65], [69], [70].

The two different frequencies used in this study provide insights into the mechanism involved in the stimulation. Three classes of mechanisms could be involved—heating, radiation force, and particle displacement. Heating is due to the absorption of ultrasound by tissues, and absorption increases with frequency. Therefore, if heating was the principal mechanism behind the stimulation, we would observe stronger effects at the higher frequency. This is not the case (Fig. 2).

Radiation force exerts pressure on a target throughout the application of ultrasound. The pressure is due to intensity gradients caused by ultrasound attenuation and reflection. Attenuation, composed of scattering and absorption, increases with frequency. Reflection, which also leads to intensity gradients, is independent of frequency. Radiation force can also be produced within standing waves. Whether radiation force could be involved in the effects reported in this study is unclear. The finding that tactile responses are produced more readily for pulsed as opposed to a continuous waveform (Fig. 5) could be suggestive of an involvement of radiation force. Yet, the finding that lower frequencies are more effective, even for pulsed stimuli, makes parsimonious explanations through a first-order approximation of radiation forces [71] difficult. Future work should measure the distribution of radiation forces in the target tissues for conclusive evidence. The responses may also result from displacements and strains of the tissue induced by radiation forces. Tissue displacement is a function of the incident force and elastic properties of the soft tissue [22], [71]. Future studies should image or calculate the displacements using careful measurements of radiation forces throughout the stimulated tissue.

We observe actions of a mechanism that is magnified at lower frequencies. On this front, the time-varying pressure wave associated with ultrasound periodically displaces particles and molecules. This cycle-by-cycle phenomenon is commonly referred to as “particle displacement”, and the displacement increases with decreasing frequency.” The maximal displacement of a particle or molecule occurs over half of the period: 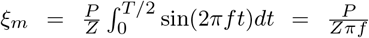. It is apparent that the displacement is inversely proportional to frequency *f*. For our 1.3 MPa, 300 kHz stimulus, the maximum displacement, using the above equation, reaches 0.92 micrometers in water. Displacements of such amplitude may be sufficient to periodically activate mechanoreceptors and ion channels [58], [72]. The effects reported in this study are unlikely due to cavitation because the lowest and medium pressures are below the mechanical index of 1.9 by the FDA [57], yet produced substantial responses (Fig. 3-top).

Our result applies to the common range of neuromodulatory frequencies, 0.25–1 MHz. At higher frequencies, especially above 10 MHz, the absorption of ultrasound by biological tissues becomes substantial [61]. This can produce appreciable intensity gradients and, consequently, radiation forces that are sufficient to activate ion channels in the peripheral nervous system [24], [25]. In addition, the high absorption can also lead to appreciable rise in temperature that can also activate ion channels even at relatively low pressures [64]. Therefore, ultrasound neuromodulation frequency likely comprises a U-shaped stimulation efficacy function. It should be noted that this proposition pertains to the peripheral nervous system. In the future, the aperture-controlled approach, such as that used in this study, should also be applied to cells in the central nervous system.

## V. Conclusion

In summary, this study controlled for stimulation volume and found that brief pulses of ultrasound at lower, 300 kHz frequencies are more effective in stimulating excitable cells and nerve fibers in human peripheral nervous system compared to higher, 900 kHz frequencies. This finding guides future choices of ultrasonic transducers for effective ultrasonic stimulation. In addition, the result suggests that stimulation of excitable cells and nerve fibers by ultrasound involves a mechanism related to cycle-by-cycle displacements of molecules or membranes by the mechanical pressure wave.

---

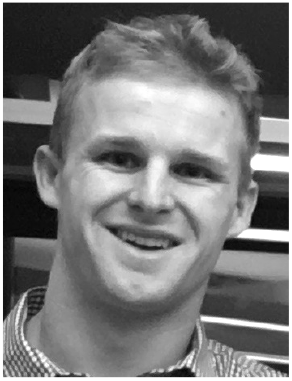

**Thomas S. Riis** Thomas S. Riis received the B.A. degree in applied mathematics and the B.A. degree in physics from University of California, Berkeley, CA, USA, in 2016. He is currently pursuing the Ph.D. degree in biomedical engineering at the University of Utah, Salt Lake City, UT, USA. He joined the Allen Institute for Brain Science in 2016, where he worked on electrophysiological recordings and optogenetic stimulation of cortex in mice. In 2018, he joined Dr. Jan Kubanek’s laboratory, University of Utah, where he works on developing new ways to modulate the nervous system noninvasively and in a targeted manner.

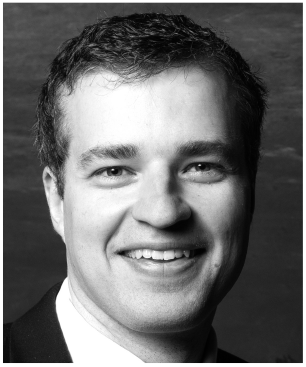

**Jan Kubanek** Jan Kubanek received his Ph.D. in Biomedical Engineering from Washington University in 2013. He obtained postdoctoral training from Stanford University, where he investigated the mechanism of ultrasonic stimulation and applied the approach to modulate choice behavior of awake behaving primates. In 2018, he joined the Department of Biomedical Engineering at the University of Utah as a tenure-track faculty. He received the K99/R00 Award from the NIH for his work. In addition, he was designated as a Moore Fellow by the University of Utah. His lab, which comprises four PhD students and one postdoc, is funded by the NIH and private foundations. The goal of the lab is to develop neuromodulation approaches that are noninvasive, focal, and act at depth, so that we can treat nervous system disorders in targeted and personalized ways without using drugs.

## Notes

This work was supported in part by the National Institutes of Health, Grant R00NS100986.

### Competing Interest Statement

The authors have declared no competing interest.

